# Somatic mutations and genome stability maintenance in clonal coral colonies

**DOI:** 10.1101/799643

**Authors:** Elora H. López, Stephen R. Palumbi

## Abstract

One challenge for multicellular organisms is maintaining genome stability in the face of mutagens across long life spans. Imperfect genome maintenance leads to mutation accumulation in somatic cells, which is associated with tumors and senescence in vertebrates. Colonial reef-building corals are often large, can live for hundreds of years, rarely develop recognizable tumors, and are thought to convert somatic cells into gamete producers, so they are a pivotal group in which to understand long-term genome maintenance. To measure rates and patterns of somatic mutations, we analyzed transcriptomes from 17-22 branches from each of four *Acropora hyacinthus* colonies, determined putative single nucleotide variants, and verified them with Sanger resequencing. Unlike for human skin carcinomas, there is no signature of mutations caused by UV damage, indicating either higher efficiency of repair than in vertebrates, or strong sunscreen protection in these shallow water tropical animals. The somatic mutation frequency per nucleotide in *A. hyacinthus* is on the same order of magnitude (10^−7^) as noncancerous human somatic cells, and accumulation of mutations with age is similar. Unlike mammals, loss of heterozygosity variants outnumber gain of heterozygosity mutations about 2:1. Although the mutation frequency is similar in mammals and corals, the preponderance of loss of heterozygosity changes and potential selection may reduce the frequency of deleterious mutations in colonial animals like corals. This may limit the deleterious effects of somatic mutations on the coral organism as well as potential offspring.

## Introduction

Patterns and mechanisms of genome stability maintenance are essential components in the evolution of long life spans in multicellular organisms. Longevity increases the potential for accumulation of mutations in cell lines throughout an organism’s lifetime, and the evolution of longevity has led to efficient DNA repair mechanisms and the segregation of somatic and germ lines in many taxa (Bythell et al. 2017). Germline segregation protects the genome stability of germ cells by reducing the number of divisions they undergo each generation, while allowing for higher mutation accumulation in somatic cells that do not contribute to the genetic makeup of the organism’s offspring (Weismann 1889). Although vertebrates and many other animal taxa display germline segregation, basal animal taxa, including flatworms, cnidarians, sponges, and some tunicates, are thought not to have a clear distinction between the two (Whitham and Slobodchikoff 1981; Radzvilavicius et al. 2016). These organisms are thought to produce gametes from pluripotent stem cells in their somatic tissues instead of harboring a protected germ line (Petralia et al. 2014). The evolutionary relationships among clonality, coloniality, and germline development are complex, but the germline may have evolved at or near the origin of the first bilaterians (Blackstone and Jasker 2003).

The role of somatic mutations in generating heterogeneity among the cells of a single organism has been theorized for decades. On the organismal level, the potential negative consequences of somatic mutation can include tumor growth, parasitism, and impaired cellular function, while potential benefits include increased genetic diversity and synergism between cells with different genetic variants (Pineda-Krch and Lehtila 2004). The empirical evidence on the cost of somatic mutations is extensive, particularly within humans and vertebrate models, because of the need to understand somatic cancer evolution to develop treatments (Pleasance et al. 2009; Forbes et al. 2015). By contrast, much of the interest in plant and germline-unsegregated animals comes from the potential for somatic mutations in these organisms to be passed down to the next generation, and thus affect the per-generation mutation rate and consequently genetic diversity on the population level (Whitham and Slobodchikoff 1981; Klekowski and Godfrey 1989; Gill et al. 1995; Orive 2001).

The role of selection on somatic mutations has been hypothesized and modeled for many taxa including plants and colonial animals, but empirically measured in far fewer taxa. When non-neutral somatic mutations occur, they can be subject to selection. Purifying selection acts strongly on germline mutations in many taxa, as deleterious mutations are purged in resulting embryos (Martincorena et al. 2015). Such selection is seen rarely in human somatic mutations, perhaps because single deleterious mutations are more permissible in a cell embedded in a functioning tissue (Martincorena et al. 2017). However, some somatic mutations erode cell-cell cooperation: in vertebrates, somatic mutations can interfere with the control of cell proliferation on which multicellular body plans depend, causing cancer. Cancerous cells tend to evolve neutrally or via positive selection, as measured by dN/dS ratios (Martincorena et al. 2017). Driver mutations are positively selected for because they are advantageous to the particular cancer cell line, but the tumor that develops is deleterious to the organism overall.

In plants, some somatic mutations appear adaptive for the cell as well as the overall organism. For example, in one study of a *Eucalyptus* tree, somatic mutations present in one set of branches produced high levels of secondary metabolites that deterred herbivory in those branches (Padovan et al. 2013). Otto and Orive (1995) developed a model which predicted that within-individual somatic selection purges deleterious mutations and thereby reduces accumulation of deleterious mutational burden in the individual. Selection on the level of individual cells, tissues, or modules of an organism may thereby contribute to the overall longevity and/or health of the organism as a whole. In addition, if the somatic mutations are heritable, then selection for beneficial somatic mutations may contribute to adaptation in the population overall (Buss 1983).

Understanding the extent to which somatic mutation rates and patterns are conserved across the tree of life may help to clarify their role in shaping organismal longevity and/or in shaping the adaptive potential of populations. Lack of knowledge about somatic mutations in diverse taxa currently hinders efforts to make generalized statements about how or why rates and patterns differ, or how those differences either drive or are driven by differences in development and life history, such as germline segregation. Far more is known about germ line and somatic cell mutations in humans and other model vertebrate systems than any other group. Human somatic base-substitutional rates are on the order of 10^−7^ to 10^−8^ per site per cell division, 3-4 orders of magnitude higher than the human base-substitutional germ line rate of 6 × 10^−11^ per site per germ cell division (Lynch 2010a). These differences in error rates are thought to stem from higher levels of DNA repair in germ cells. If the human germ line were not segregated, the number of germ cell divisions per generation would be much larger, so the heritable, per-generation mutation rate would be orders of magnitude higher (Lynch 2010b). Indeed, accounting for post-zygotic somatic mutation rates increases estimates of the per-generation mutation rate of the long-lived conifer species *Picea sitchensis* (Hanlon et al. 2019).

Many kinds of somatic mutations derive from errors in cell DNA replication during cell division (Kunkel 2004). Others derive from faulty repair of damaged DNA bases from exogenous sources such as UV. Still others come as a consequence of DNA repair, particularly repair of double-stranded breaks, during which one repair mechanism, homologous mitotic recombination, uses one strand as a template to repair the other. If the repair site had previously been heterozygous, then the heterozygous base becomes converted the homozygous state, resulting in a loss of heterozygosity (LoH) (Tischfield 1997; Tomlinson 2006). LoH mutations via gene conversion are thus distinct from gain of heterozygosity mutations that introduce a new mutant allele into the cell population. The frequency of LoH is 10^−7^ per cell division in normal human cells, but it can be as high as 10^−2^ in cells with chromosomal instability (Nowak et al. 2002). In asexual bdelloid rotifers, loss of heterozygosity as a result of gene conversion across many generations helps to explain the low heterozygosity observed in these species, and may reduce or slow down the effect of Muller’s ratchet on the population (Connallon and Clark 2010; Flot et al. 2013).

Reef-building corals comprise a highly diverse set of species of colonial cnidarians. Cnidarians have generally been considered to lack a sequestered germ line, mostly due to work on hydras that showed that stem cells could be induced to differentiate into either somatic or germ cells (Bosch and David 1987). However, followup studies have suggested that some interstitial stem cell subpopulations in hydras differentiate only into germ cells (Watanabe et al. 2009). Similarly, it is possible that Scleractinian corals possess a combination of a reservoir for germ cells and a capacity to produce germ cells from multipotent stem cells. One study has identified the *piwi* gene, a possible marker of germ line cells in many metazoans, in the coral *Euphyllia ancora* and found that *piwi*-expressing cells were present even when the polyps were not sexually reproducing (Shikina et al. 2015). The authors posited that this was evidence of germ line segregation in that coral species.

Colonial growth of an adult coral occurs through asexual budding of one polyp into two daughter polyps (Harrison 2011). Although asexual budding is expected to produce identical polyps, this is not always the case, and there are documented cases of tumor growth in corals (Coles and Seapy 1998) as well as their cnidarian relative *Hydra* (Domazet-Lošo et al. 2014). The common belief that corals do not age has come into question recently, with some evidence showing reduced telomere length with age (Tsuta et al. 2014) and other data showing lack of telomere length reduction with age for a different species (Tsuta and Hidaka 2013). Because individual genotypes can live for hundreds or thousands of years, coral biologists have made efforts to estimate the amount of variation in a single colony and what effect that might have on the adaptation rate of corals (van Oppen et al. 2011). One study indicates that intra-colony variation is common: based on microsatellite markers, intra-colony genetic variation was found in 22.8% to 46.6% of colonies (Schweinsberg et al. 2015). Estimation of somatic mutation rates across genets of *Acropora palmata* have led to age estimates up to 6,500 years among widely spread clones (Devlin-Durante et al. 2016). Another microsatellite-based study found intra-colony variability in ten of fourteen *Acropora hyacinthus* colonies investigated, and found that four of the ten colonies transferred their mutant genotypes to the eggs they spawned (Schweinsberg et al. 2014). One conflicting study found intracolony variation in the coral *Orbicella flaveolata* using the next-generation 2bRAD sequencing approach, but did not detect any of the mutations in the sperm released by the colony (Barfield et al. 2016). This study is consistent with separate germ and somatic lines in this species, and with lower levels of germ line than somatic mutation. The discrepancies between these studies drive the continued debate about the existence of germ line segregation and mutation in corals.

In this study, we measured the incidence rate and pattern of somatic mutations found in colonies of the reef-building coral species *Acropora hyacinthus*. These table top corals grow up to several meters in width over 50-200 years of growth (for review of growth rates see Gold and Palumbi 2018). This species can occur at depths of 0-25 meters (Wallace 1999), and all colonies in this sample were collected at 1-2 meters depth. We took advantage of transcriptomes sequenced from 17-22 different parts of the same colonies (Ruiz-Jones and Palumbi 2015; Ruiz-Jones and Palumbi 2017) to examine the samples for within-colony SNP variation. We verified these mutations with Sanger sequencing, measured the spectrum of somatic mutations, and documented the pattern of overall change in heterozygosity due to somatic mutations. This provides directions for further study into how these animals maintain their genomes across their long lifespans and continual growth.

## Results

### Identifying putative mutations

The first step in the mutation discovery process was to identify variation among samples taken from the same colony. We called single nucleotide polymorphisms (SNPs) across the transcriptomes of 17 different branches for each of three colonies (AH06, AH75, and AH88) and 22 different branches of one colony (AH09) (Table 1) and filtered out variants with low read counts and/or low quality scores. Only the SNPs that passed all filters and had a genotype call for every sample in the same colony were evaluated for putative mutations. We defined a putative mutation as a base with a different genotype among branches from the same colony. The mutant genotype was assumed to be the genotype least often observed in the colony (usually once), while the zygotic genotype was assumed to be the genotype most commonly seen in the colony. Rare mutations that occurred early in the growth of the colony and subsequently swept to high frequency (> 16/17 = 94%) might be recorded as having occurred in the reverse polarity under this criterion.

**Table 1.** Summary of somatic mutation identification results for *Acropora hyacinthus* colonies.

| Colony ID | Number of samples genotyped | Number of bases assayed | Colony surface area (cm<sup>2</sup>) from (Gold and Palumbi 2018) | # Putative mutations, (total #, #GoH, #LoH) | Inconclusive after Sanger | Falsified by Sanger | #Verified by Sanger (total #, #GoH, #LoH) | % putative mutations verified |
| --- | --- | --- | --- | --- | --- | --- | --- | --- |
| AH06 | 17 | 2,635,714 | 610.105 | 55, 8, 47 | 30 | 22 | 3, 1, 2 | 5.5 |
| AH09 | 22 | 5,285,809 | 13527.261 | 202, 91, 111 | 113 | 59 | 30, 11, 19 | 14.9 |
| AH75 | 17 | 6,485,057 | 1691.309 | 61, 17, 44 | 39 | 16 | 6, 1, 5 | 9.8 |
| AH88 | 17 | 5,941,437 | 382.409 | 52, 15, 37 | 22 | 18 | 12, 7, 5 | 23.1 |

**Table 2.**
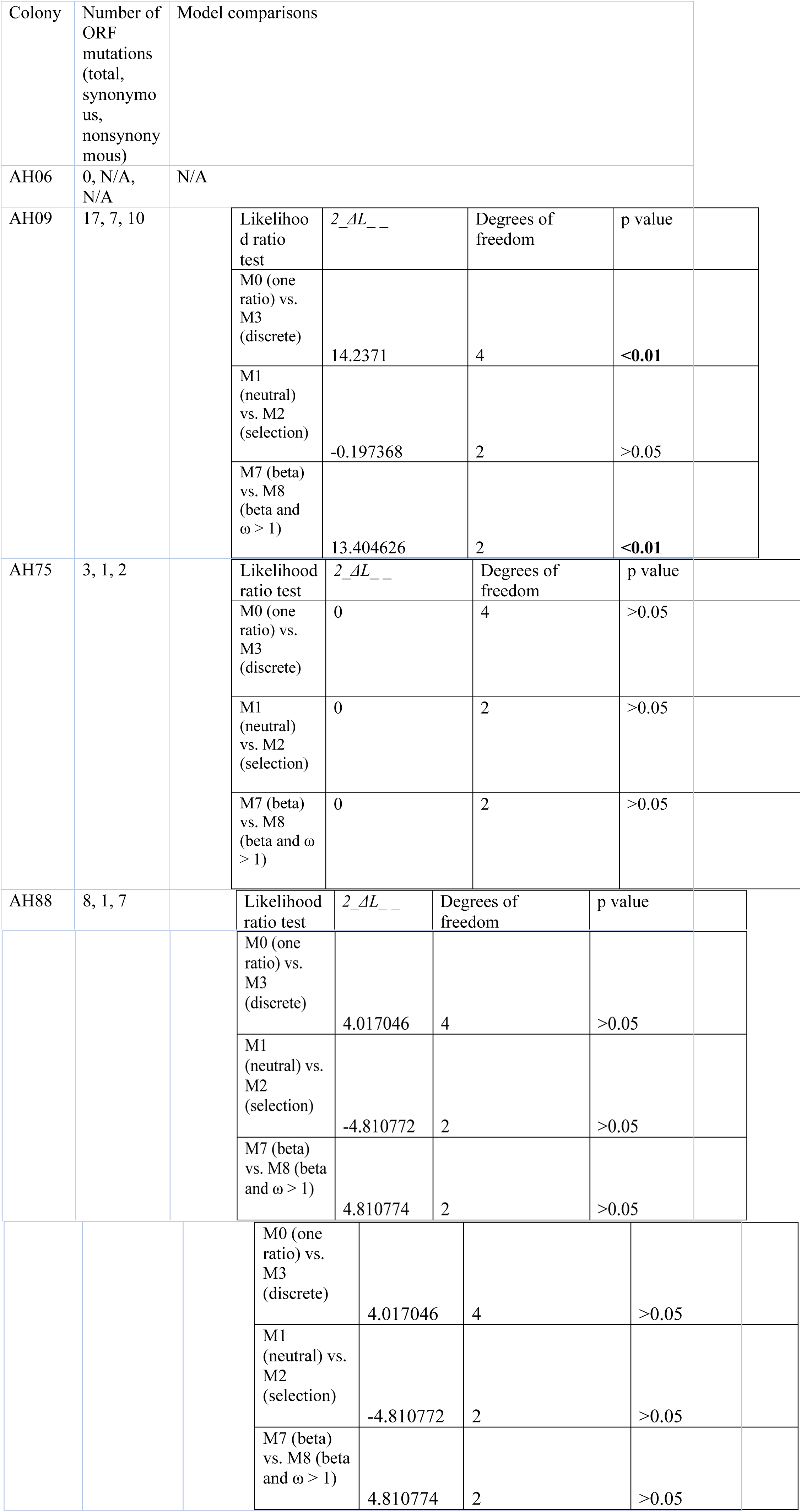
Model comparisons from likelihood ratio tests of selection within coral colonies

| Colony | Number of ORF mutations (total, synonymous, nonsynonymous) | Model comparisons |  |  |  |  |  |  |
| --- | --- | --- | --- | --- | --- | --- | --- | --- |
| AH06 | 0, N/A, N/A | N/A |  |  |  |  |  |  |
| AH09 | 17, 7, 10 | | Likelihood ratio test | $2_{-}\Delta L_{-}$ | Degrees of freedom | p value | | |
|  |  |  | M0 (one ratio) vs. M3 (discrete) | 14.2371 | 4 | <0.01 |  |  |
|  |  |  | M1 (neutral) vs. M2 (selection) | -0.197368 | 2 | >0.05 |  |  |
| | | | M7 (beta) vs. M8 (beta and $\omega > 1$) | 13.404626 | 2 | <0.01 | | |
| AH75 | 3, 1, 2 | | Likelihood ratio test | $2_{-}\Delta L_{-}$ | Degrees of freedom | p value | | |
|  |  |  | M0 (one ratio) vs. M3 (discrete) | 0 | 4 | >0.05 |  |  |
|  |  |  | M1 (neutral) vs. M2 (selection) | 0 | 2 | >0.05 |  |  |
| | | | M7 (beta) vs. M8 (beta and $\omega > 1$) | 0 | 2 | >0.05 | | |
| AH88 | 8, 1, 7 | | Likelihood ratio test | $2_{-}\Delta L_{-}$ | Degrees of freedom | p value | | |

|  |  |  |  |  |  |  |  |
| --- | --- | --- | --- | --- | --- | --- | --- |
|  |  |  | M0 (one ratio) vs. M3 (discrete) | 4.017046 | 4 | >0.05 |  |
|  |  |  | M1 (neutral) vs. M2 (selection) | -4.810772 | 2 | >0.05 |  |
| | | | M7 (beta) vs. M8 (beta and $\omega > 1$) | 4.810774 | 2 | >0.05 | |

Among 2.6 – 6.5 × 10^6^ bases sequenced per transcriptome, we identified 55 putative somatic mutations across 17 samples within colony AH06, 202 putative somatic mutations across 22 samples within AH09, 61 putative somatic mutations across 17 samples within AH75, and 52 putative somatic mutations across 17 samples within AH88 (Figure 1). Each putative mutation was additionally classified as either a loss of heterozygosity or gain of heterozygosity mutation, and as either a polymorphism seen in the population previously, or as a novel mutant never before seen in population-level studies of *Acropora hyacinthus* in Ofu, American Samoa. A list of all putative mutations, the branches that they appeared in and their status is in Supplemental Tables S1-S4. A list of all filtered, verified mutations for all colonies is in Supplemental Table S5.

**Figure 1.**
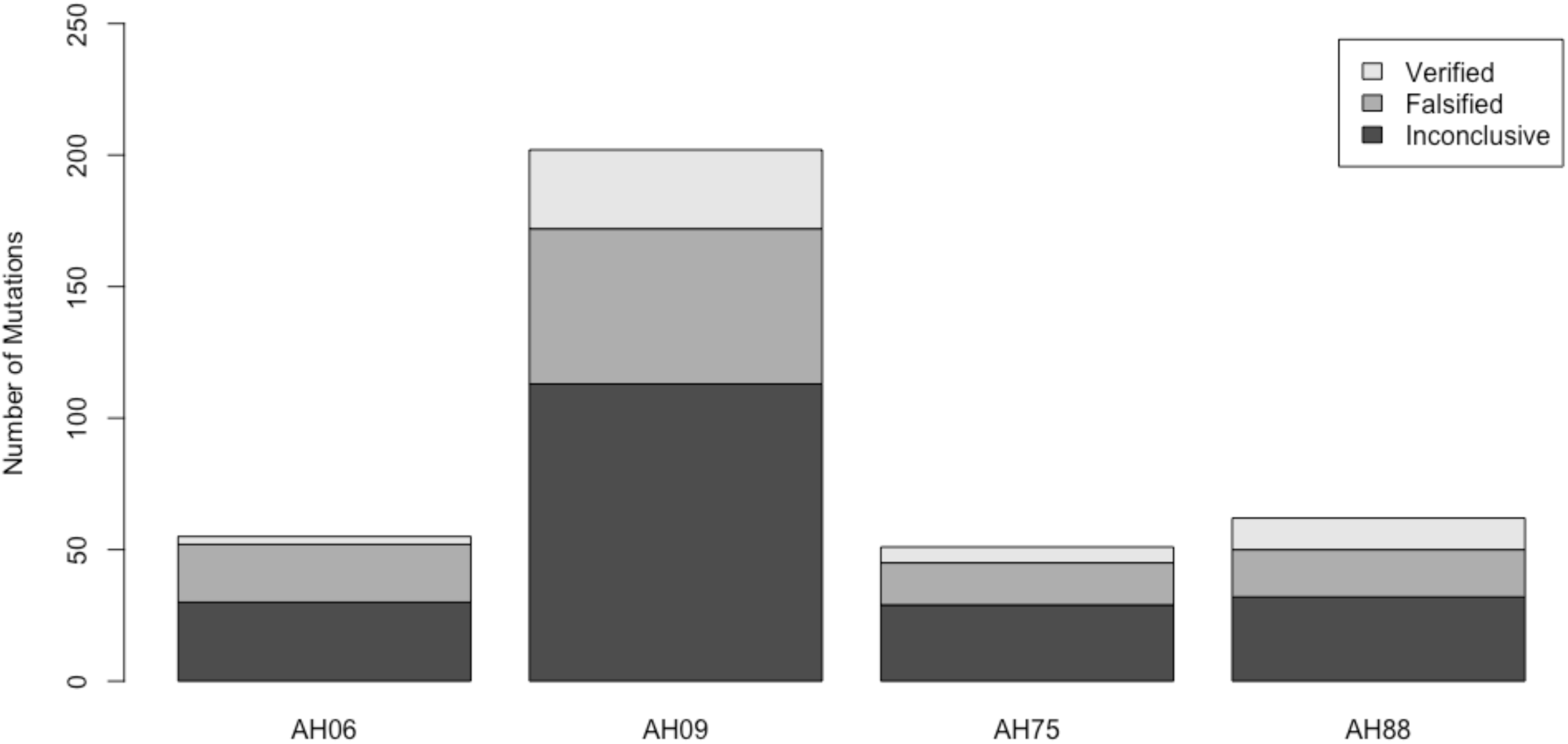
The number of mutations deemed inconclusive, false, or verified after Sanger sequencing.

### Verifying putative mutations

We then Sanger sequenced genomic DNA to verify the putative mutations for each colony. This led to a total of 3 verified somatic mutations in AH06, 30 for AH09, 6 for AH75, and 12 for AH88 (Table 1). There were no significant differences in read depth for all of the putative mutations, the falsified mutations, or the verified mutations for AH06, AH75, or AH88 alone (Supplemental figure S1a,c,d). However, there was a significant difference in mean read depth between the falsified and verified mutations for AH09 (p=0.045, Wilcoxon rank sum test) (Supplemental figure S1b). There was also a significant difference in mean read depth for between falsified and verified mutations across all colonies (p= 4.108e-05, Wilcoxon rank sum test) (Supplemental figure S1e). Neighbor-joining trees of the genetic variation within a colony did not show a pattern of relatedness among samples that corresponded to our limited information about the spatial distance among samples (Supplemental figures S2-S5). However, our power to detect such a trend is limited to the small number of mutations that occurred in more than one branch of the colony.

### Gain of heterozygosity mutations vs. loss of heterozygosity mutations

Mutations resulted in either a gain of heterozygosity (GoH) or a loss of heterozygosity (LoH). We defined gain of heterozygosity as a class of mutation in which the mutant was heterozygous but the inferred zygotic genotype was homozygous (Figure 2a). We defined loss of heterozygosity as a class of mutation in which the mutant genotype was homozygous but the zygotic genotype was heterozygous (Figure 2a). On average there were more LoH than GoH mutations in a colony (p = 0.057, Wilcoxon rank sum test). On average 36.25% +/- 17.12% (2 standard errors) of the verified mutations were GoH and 63.75% +/- 17.12% (2 standard errors) were LoH (Figure 2b), but there was considerable variation across the four colonies (Figure2c-f).

**Figure 2.**
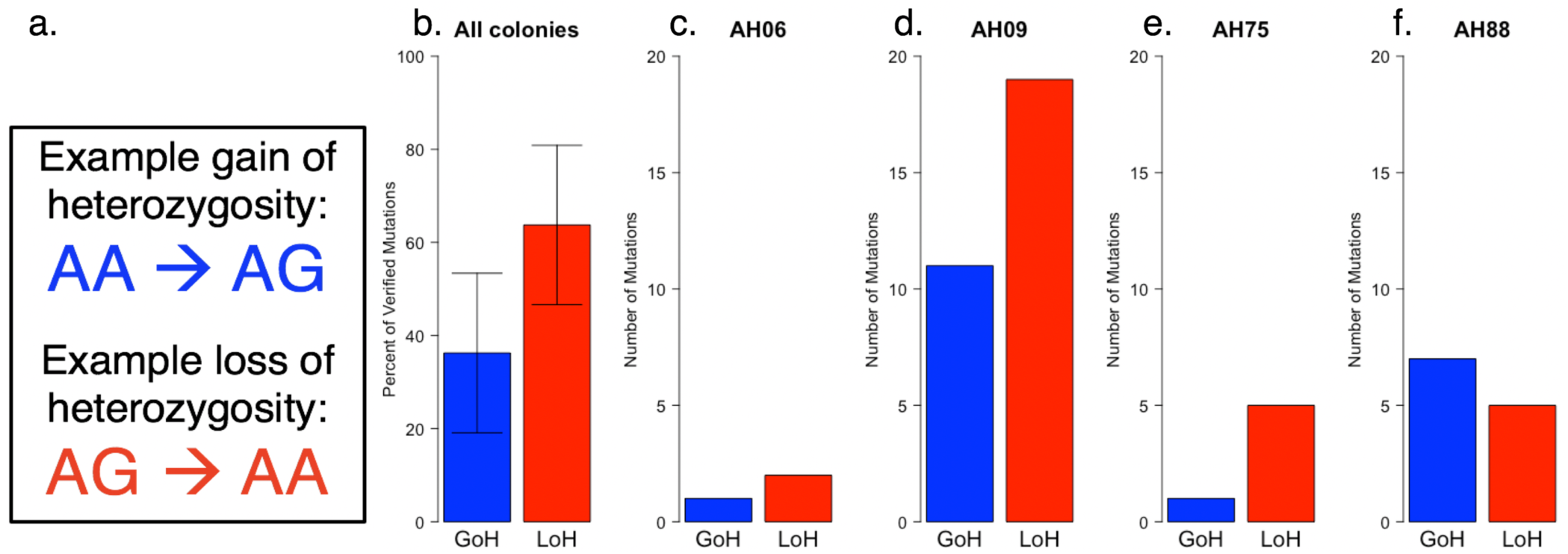
A comparison of two classes of mutation, GoH and LoH. a.) An example set of genotype changes for each class of mutation, b.) the mean percent GoH and LoH +/- 2 standard errors for all colonies, and the number of GoH and LoH mutations for c.) AH06 (N=3), d.) AH09 (N=30), e.) AH75 (N=6), and f.) AH88 (N=12).

### Population level polymorphisms vs. never seen in the population before

We compared all verified mutations against a list of SNPs that had been previously identified among 29 individual *Acropora hyacinthus* colonies in American Samoa from the same back reef pools as our study (Bay and Palumbi 2014). The majority of verified mutations (41 out of 51, 80%) had not been identified as SNPs in the population before (Supplemental figure S6). All but two of the mutations at sites that were known SNPs in the population were LoH mutations, meaning that the mutation resulted in a loss of that SNP, rather than that the mutation rose repeatedly (Supplemental figure S6).

### Relationship between mutation frequency and size

The surface area of each colony was estimated by Gold and Palumbi (2018). While size does not provide a precise estimate for age in this species, it serves as a proxy for the relative number of cell divisions occurring between existing branches, independent of absolute colony age. The frequency of mutations per nucleotide per sample was calculated by dividing the number of verified mutations by the total number of bases assayed and by the number of samples genotyped. The total number of verified mutations per nucleotide per sample was 6.7 × 10^−8^ for AH06, 2.6 × 10^−7^ for AH09, 5.4 × 10^−8^ for AH75, and 1.2 × 10^−7^ for AH88 (Figure 3). Linear regression models indicate a positive but not significant relationship between the estimated surface area of the colony and the total number of verified mutations per nucleotide per sample (p=0.07), as well as the number of verified GoH mutations (p=0.29). There was a positive and significant relationship between the colony surface area and the number of verified LoH mutations (p=0.006) per nucleotide per sample (Figure 3). This effect is driven by the higher frequency of mutations seen in AH09, the largest colony, versus the three smaller colonies.

**Figure 3.**
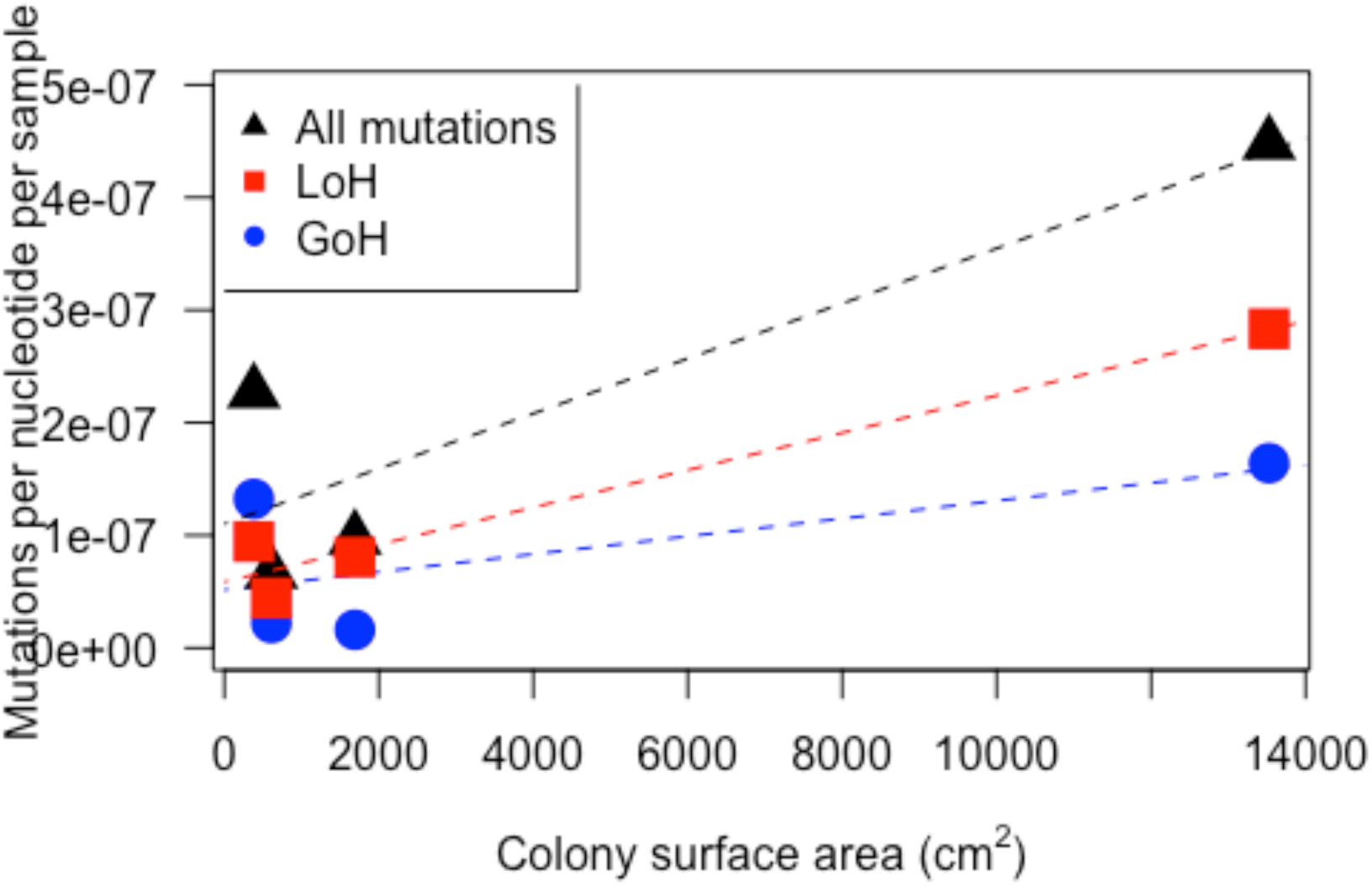
The number of gain of heterozygosity (GoH), loss of heterozygosity (LoH), and total mutations per nucleotide per sample in each colony scales positively with colony size.

### Mutational spectra

Most of the individual colonies in this study had very few mutations, so the spectrum of each one looks quite different from the others due to low sample size (Supplemental Figure S7). A clearer pattern resolves when the mutations from all four colonies are combined (Figure 4a). The mutation spectrum of all verified mutations from all colonies is not significantly different from the mutation spectrum for just the verified LoH mutations (chi-square test, p=0.76) or the mutation spectrum for just the verified GoH mutations (chi-square test, p=0.58) (Supplemental Figure S8). The transition/transversion ratio of the combined set of all verified mutations in the study is 1.68.

**Figure 4.**
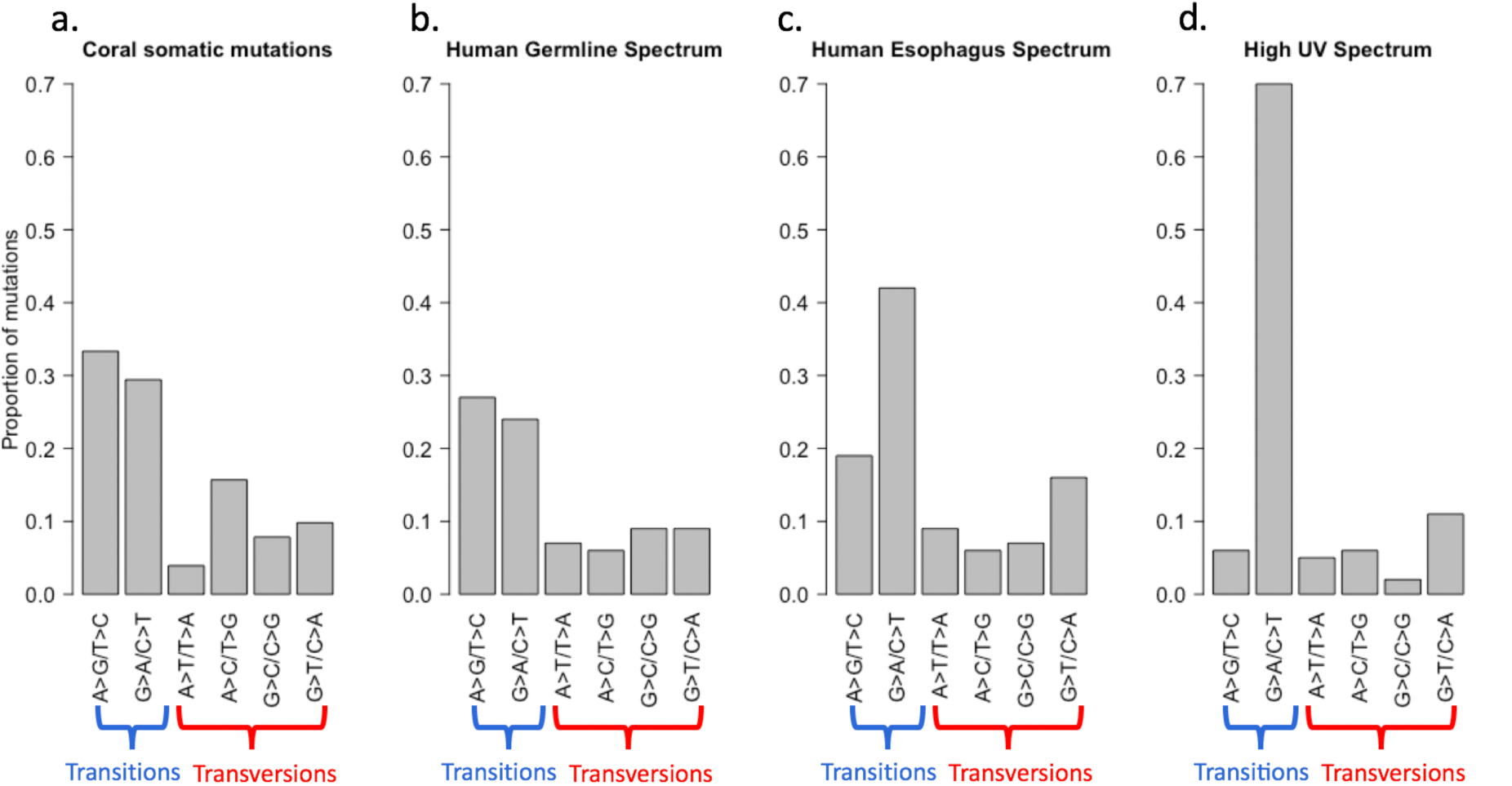
Mutation spectra for somatic mutations from different types of tissues. a.) the verified somatic mutations for the four *Acropora hyacinthus* colonies in this study (N=51) b.) human germline mutations (Rahbari et al. 2016), c. human somatic mutations in esophagus cells (Martincorena et al. 2018), and d.) UV-induced mutations in human melanoma (Pleasance et al. 2009).

We compared the coral somatic mutation spectrum (Figure 4a) with the spectrum for human germline mutations (Rahbari et al. 2016) (Figure 4b), the spectrum for human esophagus somatic mutations (Martincorena et al. 2018) (Figure 4c), and the spectrum for human melanoma somatic mutations (Pleasance et al. 2009) (Figure 4d). The same signature in human melanoma is also seen in non-cancerous, sun-exposed human skin cells (Martincorena and Campbell 2015). The coral somatic mutation spectrum is not significantly different from the human germline mutation spectrum (chi-square test, p=0.22), but it is significantly different from the human esophagus somatic mutation spectrum (chi-square test, p=2.0 × 10^−11^) and the UV damage-induced melanoma mutation signature (chi-square test, p= 1.7 × 10^−15^). This is striking, considering the high UV environment these corals inhabit (Shick et al. 1996).

### Selection on mutations as monitored by amino acid sequence changes

Tests of selection yielded different results for each colony, likely driven by low numbers of coding mutations. None of the three verified AH06 mutations were in the open reading frame so dN/dS could not be calculated. The dN/dS values are equal to or greater than 1 for all models for AH88 (Supplemental Table S6), but less than or around equal to 1 for AH09 (Supplemental table S7) and AH75 (Supplemental Table S8). However, no comparison rejected the null hypothesis of neutral evolution (M1) for any colony. Six sites were found to be under significant positive selection in AH88 under M2, M3, and M8 (Supplemental Table S6) and one site was found to be under significant positive selection in AH09 under M3 and M8 (Supplemental Table S7).

### Linked changes

We found two instances in which multiple verified mutations were present within the same contig. In each instance the mutated state was seen in only one sample among all samples genotyped. One example of such closely occurring mutations, observed in one sample in colony AH09, consisted of three LoH mutations that occurred across a span of 12 nucleotides (Figure 5a). The other example of closely occurring mutations, observed in one sample in colony AH88, included one LoH and three GoH mutations that occurred across a span of 97 nucleotides (Figure 5b). All of the mutations in each of the sets of mutations occurring on the same contig were nonsynonymous substitutions. None of these SNPs had been seen in the population previously.

**Figure 5.**
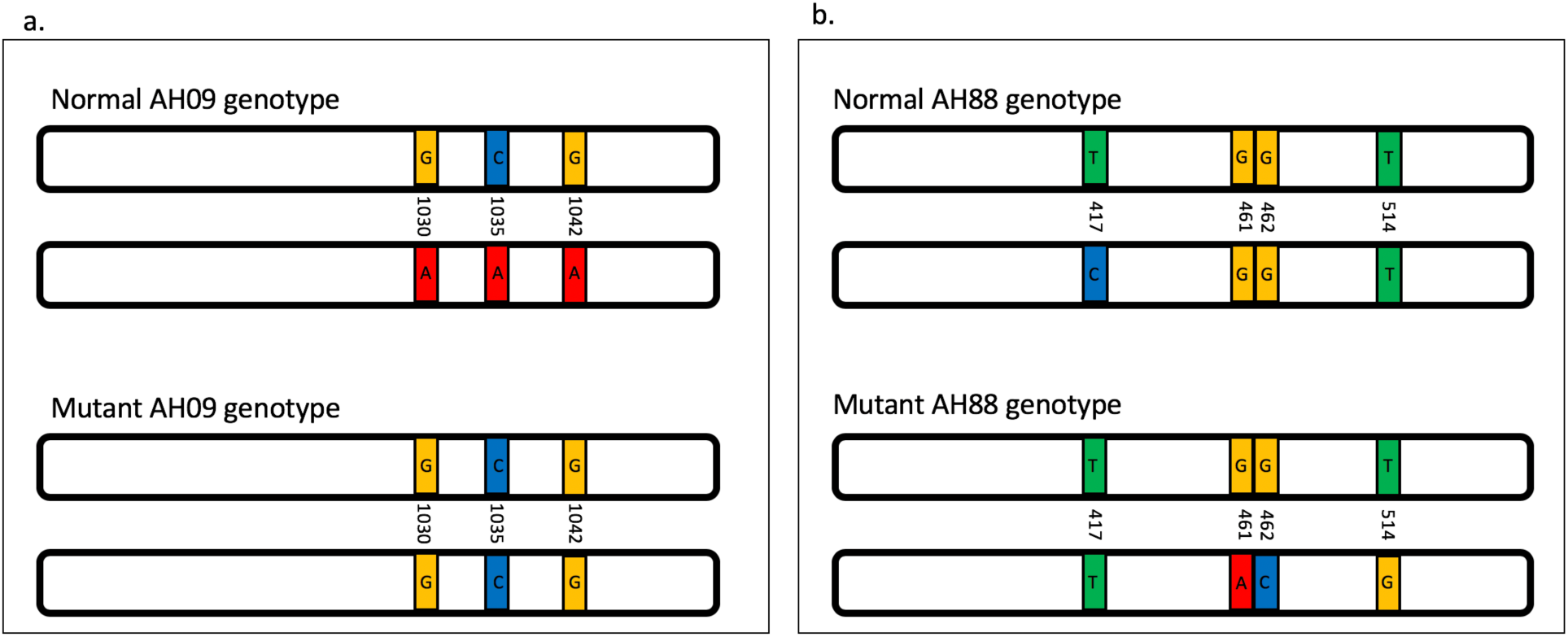
Schematic of mutations occurring in the same contig in the diploid coral samples. a.) colony AH09, in which all 3 mutations resulted in loss of heterozygosity, and b.) colony AH88, in which the mutation at position 417 resulted in loss of heterozygosity and the mutations at 461, 462, and 514 resulted in a gain of heterozygosity.

## Discussion

### Verified mutations

Of the putative mutants identified by Illumina HiSeq sequencing, 5.5-23.1% per colony were verified by Sanger sequencing of genomic DNA from the same coral sample (Table 1). This is in line with the rate of Sanger verification found in putative somatic mutations of spruce trees (Hanlon et al. 2019). Falsified mutations were likely a result of Illumina sequencing error or the differential transcription or differential sequencing of the two alleles in the transcriptome libraries. Some gain of heterozygosity mutations may have been common enough to be visible to next generation sequencing methods (occurring in ca. 10% of reads), but rare enough in a tissue sample to not be clearly visible in a sequencing chromatogram.

Even more putative mutations were deemed inconclusive, either because they would not amplify with the primers we designed for them or because the Sanger sequence chromatograms produced from these amplifications were too noisy to make a confident genotype call. There were no significant differences in read depth for putative mutations that were falsified, those that were inconclusive, or those that were verified within a colony. However, when data from all colonies were combined it emerged that verified mutations had a significantly higher read depth than falsified mutations (Supplemental Figure S1e).

### Mutation spectra

We compared the coral somatic mutation spectrum to those found in human somatic cells and those found in human germ cells in an attempt to ascertain whether the selection pressures or mechanisms of DNA repair in corals might be more similar to established patterns for soma or for germ lines. The mutation spectrum of all of the verified coral mutations shows the highest similarity with human germ line mutations: both spectra show that rates of the mutation G>A/C>T and the reverse A>G/T>C are similar. By contrast, human somatic mutations have far higher G>A/C>T changes than the reverse. This is particularly true for the mutational spectrum for UV damage in somatic tissue, which is exceptionally dominated by G>A/C>T changes Tropical, reef-dwelling corals, particularly those living in shallow (1-2 m deep) water such as the corals in this study, live in very highly UV-exposed environments (Fleischmann 1989; Shick et al. 1996). However, the luminescent skeletons of Scleractinian corals have been shown to protect the tissue from UV radiation damage by absorbing UV radiation and emitting yellow light (Reef et al. 2009). There may be other mechanisms, such as enhanced DNA damage repair mechanisms or somatic selection, that explain the lack of UV-induced mutations in these corals. Somatic mutation frequency decreases with depth in *Orbicella* coral colonies, however the authors argued that this relationship was not due to increasing UV attenuation with depth, due to the fact that the mutations they identified did not show the mutational signature characteristic of UV-induced mutations (Olsen et al. 2019).

### Selection on somatic mutations

Two different patterns emerged in our dataset: colony AH88 appeared to be dominated by positive selection, with six different sites showing evidence for significant positive selection, whereas the other colonies did not show significant positive selection. Overall many of the models produced dN/dS values lower than 1 for AH09 and AH75, suggesting that purifying selection may be acting overall on these colonies, however there was little statistical power for these tests due to small sample size. Stronger selection on coral somatic cells than human somatic cells might occur because a larger proportion of the genome is exposed to selection in the less specialized tissues of corals. Alternatively, it could result from the difference between a modular, colonial animal that has each individual polyp exposed to the environment, as opposed to a unitary animal whose cells are embedded in many tissues. It may also be a reflection of the modular structure in which cell lineages may develop into many cell types during the long lifetime of the colony. However, somatic mutations in colony AH88 were dominated by amino acid changes, with six different sites showing evidence for significant positive selection. Small sample size in our study limits our ability to finely analyze patterns of selection on mutations, and a larger genomic data set will be needed to test for any purification of deleterious mutations or positive selection on advantageous mutations in coral cell lines.

### Loss of heterozygosity over time in older corals

LoH usually results from repair of double stranded breaks in DNA. When a break occurs, one mechanism of repair of the missing or damaged DNA sequence is accomplished by homologous recombination, which copies a template from a sister chromatid or homologous chromosome and pastes it into the damaged site (Takata et al. 1998; Helleday et al. 2007). This can result in a long section of heterozygous sites being turned homozygous, as we see in colony AH09, where three sites within 12 bp of each other are heterozygous for 21 of 22 branches but all three sites are homozygous in one branch (Figure 5a). More research should be done to determine whether homologous recombination is the predominant mechanism of double stranded break repair. Additionally, rates of LoH may be different in coding and noncoding regions of the genome, so follow-up studies with full genome scans for somatic mutations will elucidate the role of homologous recombination on the genomic level.

LoH has not been the subject of focus in most theoretical literature that deals with the consequences of somatic mutations for the organism or population. We find that each of our colonies is heterogenous, but counterintuitively, reduced heterozygosity is the most common form of heterogeneity. While Buss (1993), Klekowski and Godfrey (1989), and Orive (2001) all considered the effects of somatic mutation on increased genetic diversity, none of them modeled the effects of decreased diversity, particularly on the next generation of gametes. We find that in *Acropora hyacinthus*, for every appearance of a new heterozygous site, two heterozygous sites are lost. Over the course of the coral’s life span, then, individual polyps should show a net loss of heterozygosity. However, overall heterozygosity across the colony is not lost: different cell lines within the colony together hold the original diversity seen in the zygote. There could be a long-term pressure on corals to retain sexual reproduction and recombination as the means of maintaining genetic variation within a population as a correction to the LoH that individual colonies experience. Hartfield et al. (2016) predicted using coalescence models that when sex is rare in a lineage, gene conversion becomes a significant force in reducing diversity within individuals. The bdelloid rotifer provides empirical evidence of an asexual lineage that has lost heterozygosity due to gene conversion (Flot et al. 2013). Maintaining sex in clonal, colonial animals may therefore be crucial to counteract LoH over generations.

### Somatic mutations and longevity

Some somatic mutations have been shown to scale with age, leading to genome mosaicism in an individual’s lifetime (Vijg 2014). Somatic mutation accumulation is in fact often considered a driver of aging (Kirkwood 2002). A linear relationship between age and number of somatic mutations has been found in human esophagus, brain frontal cortex, kidney cortex, and colon epithelial cells (Hoang et al. 2016; Martincorena et al. 2018). In our data, the frequency of somatic mutations per sample increased with coral colony size, dominated by higher mutation frequency in AH09, a multi-plate colony that is much larger than colonies AH06, AH75 and AH88. Growth records of AH06 and AH75 show steady increase in colony size over five years from 2010 through 2014, and these colonies were at least six years old when the samples used in this study were taken [24]. AH09 was 194 cm across when first measured in 2010 and its age is uncertain (Gold and Palumbi 2018). Growth rates of conspecifics on the same reef as these colonies suggest area growth of 5% and radial extension of about 1 cm per year (Gold and Palumbi 2018). By that metric, AH09 is on the order of 90-100 years old. Because of partial mortality, disturbance and regrowth, coral size and age are not well correlated (Gold and Palumbi 2018), this is simply an order-of-magnitude estimate.

If we use our rough estimates of the ages of the colonies in our study, coral genomes accumulate between 2.6 and 6.7 mutations per gigabase per year in coding regions: (averaging 4.9 per gigabase per year; 6.7, 2.6, and 5.4 per gigabase per year for AH06, AH09 and AH75, Supplemental Table S9). For types of mutations that accumulate with time, human somatic cell lines also accumulate 1-25 mutations per gigabase per year; tissues with quick cell turnover such as stomach and digestive linings are at the high end of that range (Alexandrov et al. 2015).

Although coral mutation frequency is similar to humans, the mutations accumulated in the coral colony may be less deleterious to the overall colony. It may be that mutant sites are filtered by selection, so that perhaps the most deleterious changes are purged in growing somatic cell lines, and advantageous mutations proliferate. This is consistent with the predictions made by Buss (1983) and Otto and Orive (1995): that intra-organismal selection will reduce the frequency of deleterious mutations in asexually reproducing organisms with plastic development that give rise to multicellular offspring. Perhaps the lack of germ line drives the necessity for heightened somatic selection (see section on the question of coral germ lines below), and therefore the differences in apparent aging between corals and humans has less to do with being colonial or unitary and more to do with the necessity to purge deleterious mutations if they have a chance of being inherited by offspring. *Acropora hyacinthus* is not an exceptionally long-lived coral compared to some other corals, so somatic mutation identification of extremely old corals will be useful to help resolve the questions of how and if corals age.

Mutation rate comparisons are more clearly resolved when looking at the mutations per base pair per mitosis rate than when looking at the mutational frequency (mutations per base pair). For instance, the mutations per base pair in the mouse germline and soma are similar to mutations per base pair in the human germline (Milholland et al. 2017). However when the mutations per base pair per mitosis is accounted for, the mouse germline and somatic mutation rates are significantly higher than the human rates (Milholland et al. 2017). This is because human cells acquire fewer mutations per mitosis than do mouse cells but have more mitoses. Higher per-mitosis mutation rate in mice than humans have been proposed as an explanation for differences in the lifespan of the two species (Milholland et al. 2017). Lack of information about cell division rates proves to be a consistent problem in estimating mutations rates of many non-model organisms, including spruce trees (Hanlon et al. 2019), and another long-lived clonal organism, the fungus *Armillaria gallica* (Anderson et al. 2018). More work on rates of cell division in coral adults, in addition to sampling strategies that explicitly take distance and positioning between samples into account, will determine whether corals have a low rate of mutation per cell division.

### Soma and germ line segregation in corals

Corals have classically been thought to convert somatic cells into germ cells – with each polyp producing gametes from their own somatic tissues. Recently, Barfield et al. (2016) found somatic variation in adult polyps but could not find those variants in the sperm produced from different patches of the same colony, and argued that somatic tissues do not differentiate into gamete producers as has been long thought. In contrast, Schweinsberg et al. (2014) claimed that they did find transfer of somatic mutations from adult polyps to their gametes for some, but not all, *Acropora hyacinthus* colonies studied. Our study does not directly resolve the question of somatic versus germ line segregation in corals or test for the existence of germ stem cell lines in these animals. However, we find numerous, verifiable mutational differences among branches of even small table corals based on moderate sequencing effort. Moreover, we see some evidence of selection, so that perhaps the most deleterious changes are purged before germ cell production. Thus, rates of potentially deleterious mutation may be low enough to allow corals to successfully produce gametes from somatic cells. This is in line with predictions made by Orive (2001) when somatic selection is accounted for in a model: that somatic selection would act against deleterious mutants in the individual and therefore the gametes that the organism produces will not carry many deleterious mutations that arose in the parent’s somatic tissues. Direct comparison of polyp and gamete genomes from the same polyps, as well as more research on stem cells and the process of reproduction in corals are needed to resolve the question of germ line segregation in corals.

### Conclusions

Our data detail patterns of mutations in somatic cells in long lived corals and allow comparison to well-known patterns in human germlines, somatic cells, and tumors. Corals accumulate somatic mutations with colony size, similar to accumulation of somatic mutations with age in humans. They also show mutation spectra more similar to highly protected human germlines than to human somatic mutations. In particular, the mutation spectrum associated with human somatic UV damage is not observed in corals, despite the shallow water exposure of corals to high UV light levels (Shick et al. 1996). Corals show more LoH than GoH mutations. A consequence of this is that polyp lineages experience net loss rather than gain of heterozygosity as the colony grows. Total mutational frequency (mutations per nucleotide in a given sample) in our largest coral is similar to that found in humans (Martincorena et al. 2018). However, selection might be stronger in corals, possibly resulting in lower levels of deleterious mutations in active cell lines. This may help to explain how long lived coral somatic tissue might give rise to gametes without showing adverse consequences.

RNA sequence data have become more and more valuable as tracers of cell lineages (Yizhak et al. 2019), as with our data, the total number of mutations per sample is small. Comparisons of full genomes across coral colonies, especially large, long lived colonies, will allow more powerful tests of the mutation and DNA repair processes that distinguish corals from better known vertebrate systems. In addition, somatic mutation rate and pattern estimates from taxa spanning more phyla in the tree of life will be necessary to determine the conservation and/or diversification of somatic mutations across different body plans, life history strategies, and evolutionary relationships.

## Materials and Methods

### Sample collection, library preparation, genotype calling

Seventeen sample were taken from each of three *Acropora hyacinthus* colonies (described in Ruiz-Jones and Palumbi (2017)) and 22 sample were taken from a fourth *Acropora hyacinthus* colony (described in Ruiz-Jones and Palumbi (2015)). For colonies AH06, AH75, and AH88, each sample was taken in a counterclockwise along the perimeter of the colony, 1-3 cm apart (Ruiz-Jones and Palumbi 2017). For colony AH09, samples were taken in pairs, with two sample adjacent to each other, but the positioning of each pair relative to the other pairs across the colony are not known. All of these colonies live in shallow, back-reef lagoons in Ofu, American Samoa (14.18206°S, 169.65849°W). The surface area of each of these colonies was calculated with ImageJ analysis by Gold and Palumbi (2018). RNA was extracted from 3-5 polyps from one branch for each sample and transcriptome libraries were prepared and sequenced (Ruiz-Jones and Palumbi 2015; Ruiz-Jones and Palumbi 2017). Reads that passed quality checks (duplicate reads removed, length of >20 base pairs, and quality score of >20) were mapped with Bowtie 2 (version 2.2.3) (Langmead and Salzberg 2012) against a reference *Acropora hyacinthus* transcriptome (Barshis et al. 2013). We called single nucleotide polymorphisms (SNPs) using the Haplotypecaller and GenotypeGVCFs tools from the Genome Analysis Toolkit version 3.7 (McKenna et al. 2010). Variants were filtered with bcftools and vcftools to include only biallelic SNPs with Phred quality score of 30 or higher. We calculated the total number of non-variant sites that met all of the same criteria as the SNPs by using the GenotypeGVCFs tool with the output mode EMIT_ALL_SITES.

### Post-Genotype Calling Analyses

We filtered SNPs to include only those that were polymorphic among branches of an individual coral colony. These were deemed putative somatic mutations. SNPs that had been previously found at the population level in *Acropora hyacinthus* colonies in Ofu (Bay and Palumbi 2014) were labeled “population polymorphisms.” We classified each putative somatic mutation as a transition or transversion. Directionality was determined by whether a genotype was seen in the minority or majority of samples in a colony. The minority genotype in a colony was always classified as the mutant state: usually the mutant was seen in only one branch (see Supplemental Tables S1-S4). Similarly, mutations were classified as generating a gain of heterozygosity (GoH) or a loss of heterozygosity (LoH). GoH mutations were defined as those where the majority of samples from a single colony were homozygous (with 0% minor allele frequency at that locus), and the minority were heterozygous (>10% minor allele frequency). LoH mutations were defined as those where the majority of samples in a colony were heterozygous at a certain position (>10% minor allele frequency), and the minority were either homozygous reference or homozygous alternate at that position (0% minor allele frequency).

### Verification

We extracted genomic DNA from the branches that provided RNA for the transcriptome libraries using Nucleospin Tissue columns and the corresponding protocol for extraction from animal tissue (Macherey-Nagel, Düren, Germany). We designed primers that sequenced 120-140 bp regions of gDNA that contained each putative somatic mutation SNP using Primer3 (Untergasser et al. 2012) and a modified GTSeq primer design pipeline (Campbell et al. 2015). PCR products were sequenced by Elim Biopharm (Hayward, CA, USA). We mapped the Sanger sequences to the relevant contigs from the *Acropora hyacinthus* transcriptome and called genotypes using Geneious 11.0.5 (https://www.geneious.com). We classified each putative mutant as verified if the Sanger electropherogram matched the Illumina genotype call or falsified if the Sanger electropherogram did not match the Illumina genotype call. If the electropherogram did not present a clear genotype call, the targeted SNP failed to amplify, or the targeted SNP amplification failed to sequence, the putative mutation was ruled inconclusive.

### Tests for amino acid sequence changes and selection

We predicted the open reading frame (ORF) for each mutation-containing contig using ORFpredictor (Min et al. 2005). We then determined whether each mutation was located in the ORF, or outside of it. We concatenated all of the mutation-containing contigs for each sample and built an alignment and a Newick tree of the samples from each colony (Supplemental Figures S1-4) using Geneious version 11.0.5. We determined maximum likelihood estimates for the ratio (w) of number of synonymous substitutions per synonymous site (d_S_) to the number of non-synonymous mutations per non-synonymous site (d_N_) using the program CODEML in the package PAML (Yang 1997). We conducted likelihood ratio tests to infer the presence/absence of selection. The likelihood statistic is 2Δl = 2(l1 – l0), where l0 is the likelihood under the null model, l1 is the likelihood under the alternative model and x2v is the distribution, where *v* is the number of degrees of freedom and corresponds with the number of free parameters. In CODEML, model M0 assumes a single value of w at all sites and lineages, whereas M3 contains a discrete distribution of values for w. M1 assumes two site classes, one under purifying selection and the other under neutral evolution, and was compared to M2, which allows for three site classes, including positive selection. We also compared M7, which assumes that variation in w has a beta distribution, to M8, which allows for positive selection.

## Acknowledgements

This work was supported by the Gordon and Betty Moore Foundation. EHL was funded by the NSF Graduate Research Fellowship and the Morgridge Family Fellowship (Stanford Graduate Fellowship in Science and Engineering). We thank L. Ruiz-Jones for collection and preparation of the samples analyzed in this study. Thanks to N. Rose and B. Barney for assistance with developing bioinformatic pipelines.

