## SupplementalFIgures for "Somatic mutations and genome stability maintenance in clonal coral colonies"

Supplemental Figures

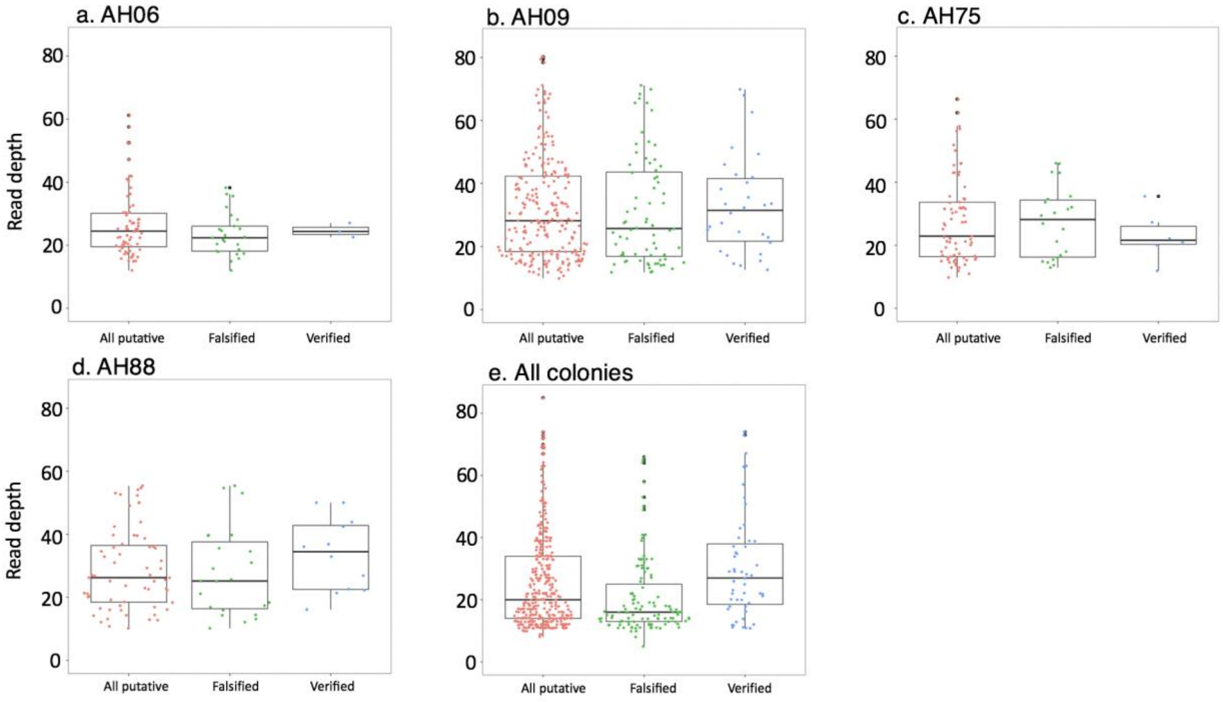


Supplemental Figure S1. Read depths for all putative, mutations, falsified mutations, and verified mutations. Data shown for a.) AH06, b.) AH09, c.) AH75, d.) AH88, e.) all colony data combined.

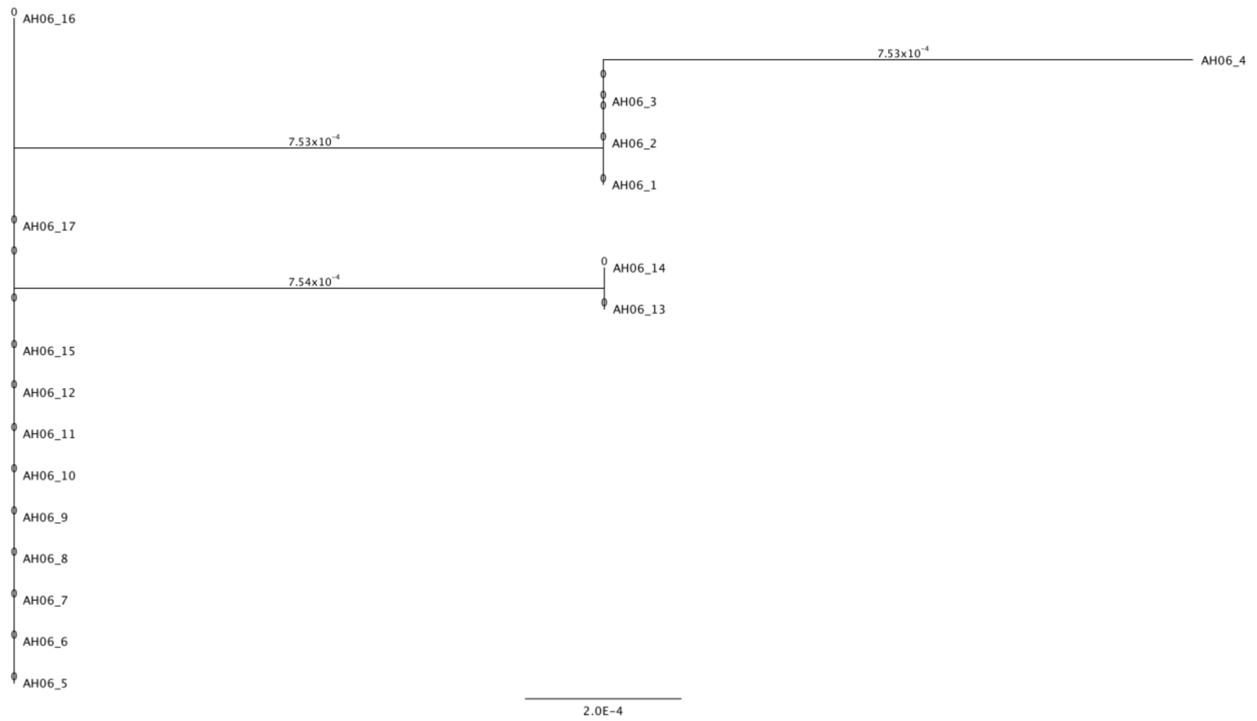


Supplemental Figure S2. Neighbor-joining genetic distance tree for colony AH06. The samples for this colony were taken sequentially in a line, such that difference between the numbers following the prefix “AH06_” corresponds with the relative spatial distance between the samples in the colony.

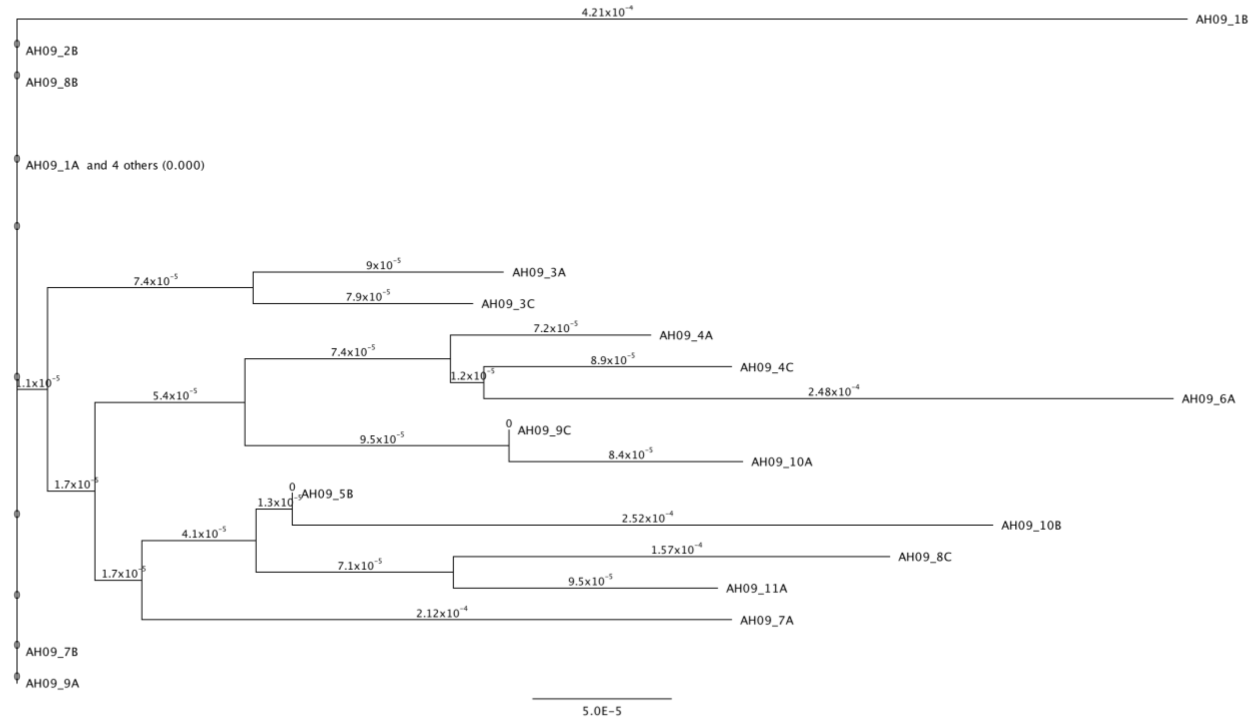

Supplemental figure S3. Neighbor-joining genetic distance tree for colony AH09. The samples from this colony were taken in pairs, such that the samples with the same number following the prefix “AH09_” were the most spatially close to one another.

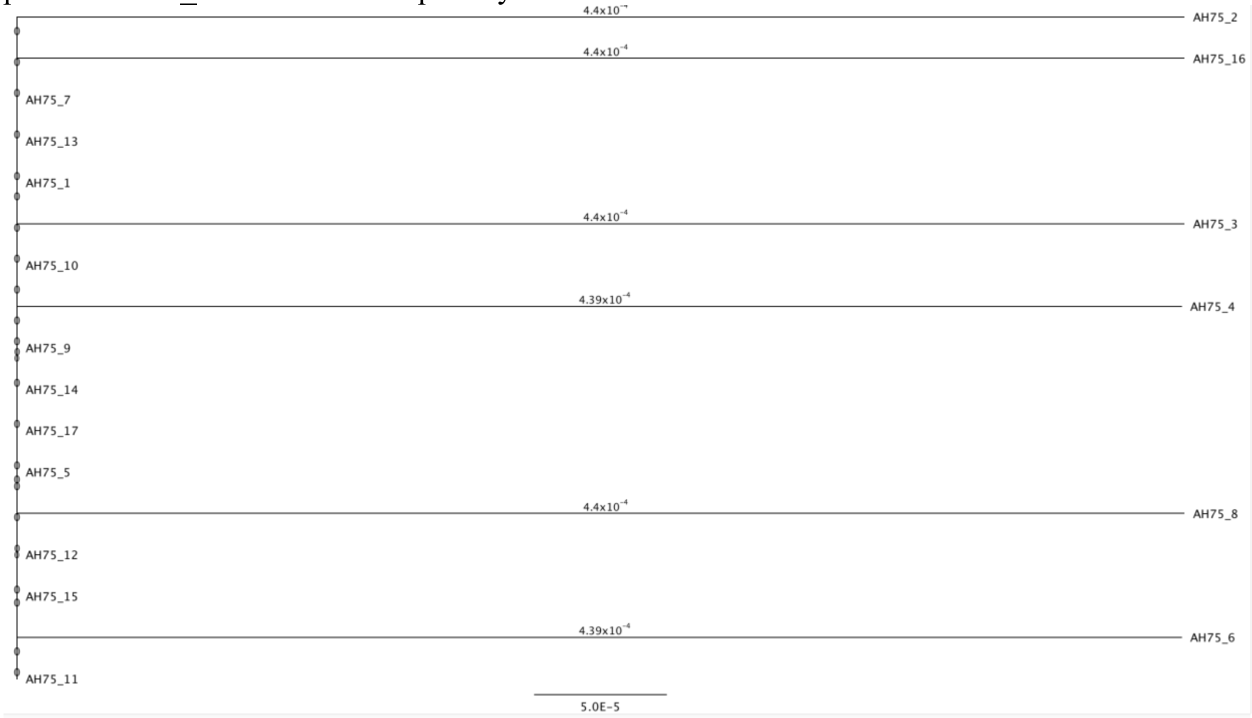


Supplemental figure S4. Neighbor-joining genetic distance tree for colony AH75. The samples for this colony were taken sequentially in a line, such that difference between the numbers following the prefix “AH75_” corresponds with the relative spatial distance between the samples in the colony.

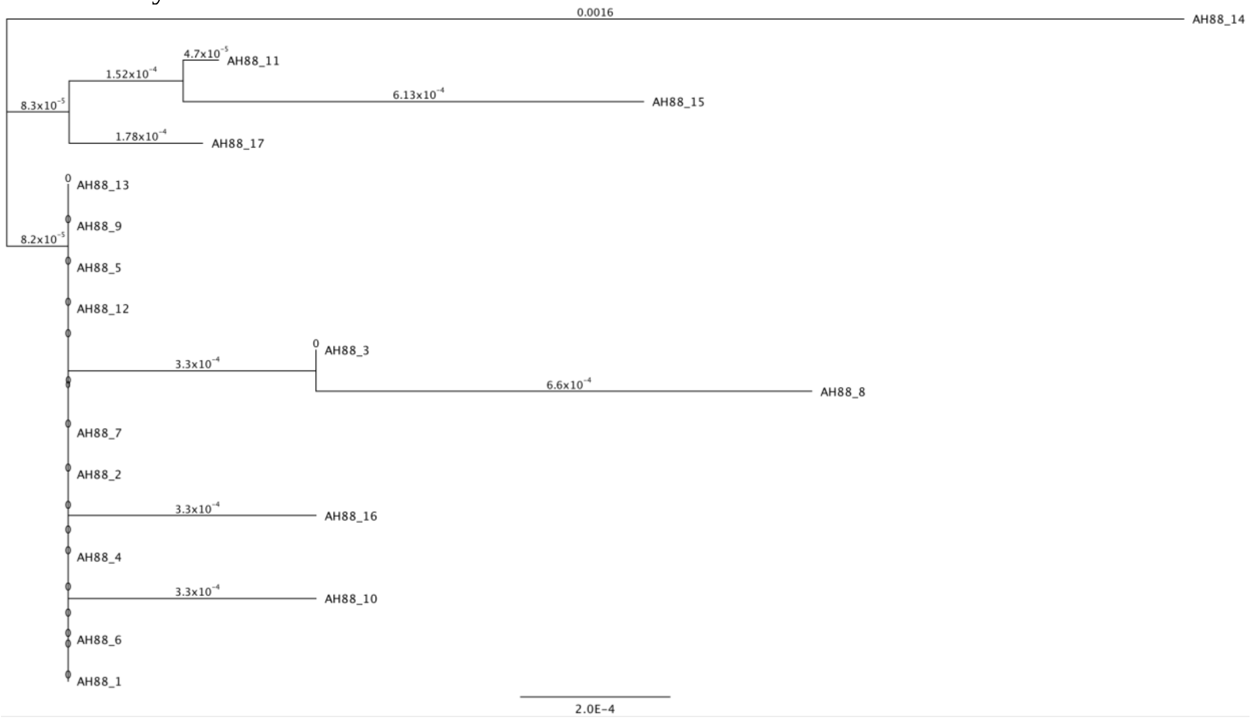

Supplemental figure S5. Neighbor-joining genetic distance tree for colony AH88. The samples for this colony were taken sequentially in a line, such that difference between the numbers following the prefix "AH88_" corresponds with the relative spatial distance between the samples in the colony.

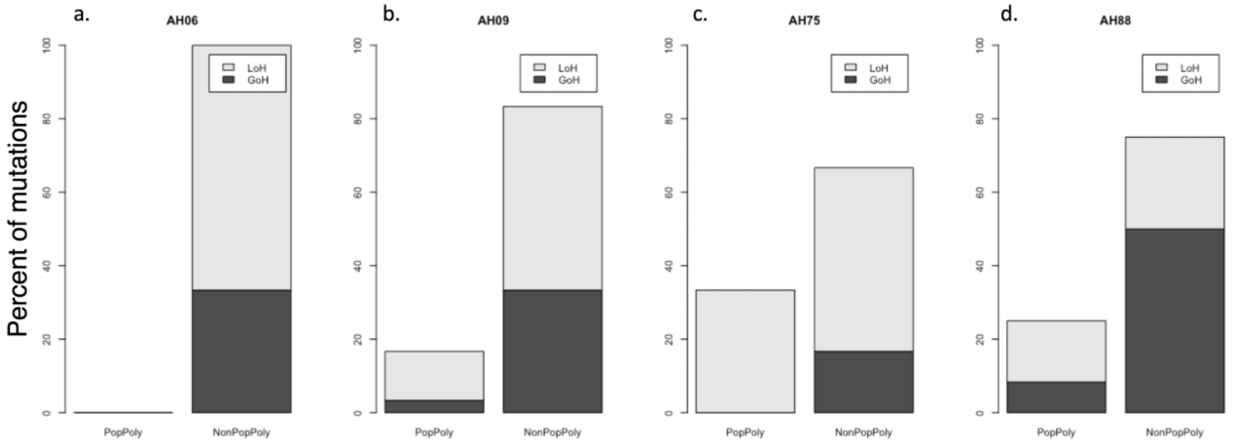


Figure S6. The percentage verified mutations that were pre-existing polymorphisms and LoH, pre-existing polymorphisms and GoH, novel mutants and LoH, and novel mutants and GoH for a.) AH06 (N=3), b.) AH09 (N=30), c.) AH75 (N=6), and d.) AH88 (N=12).

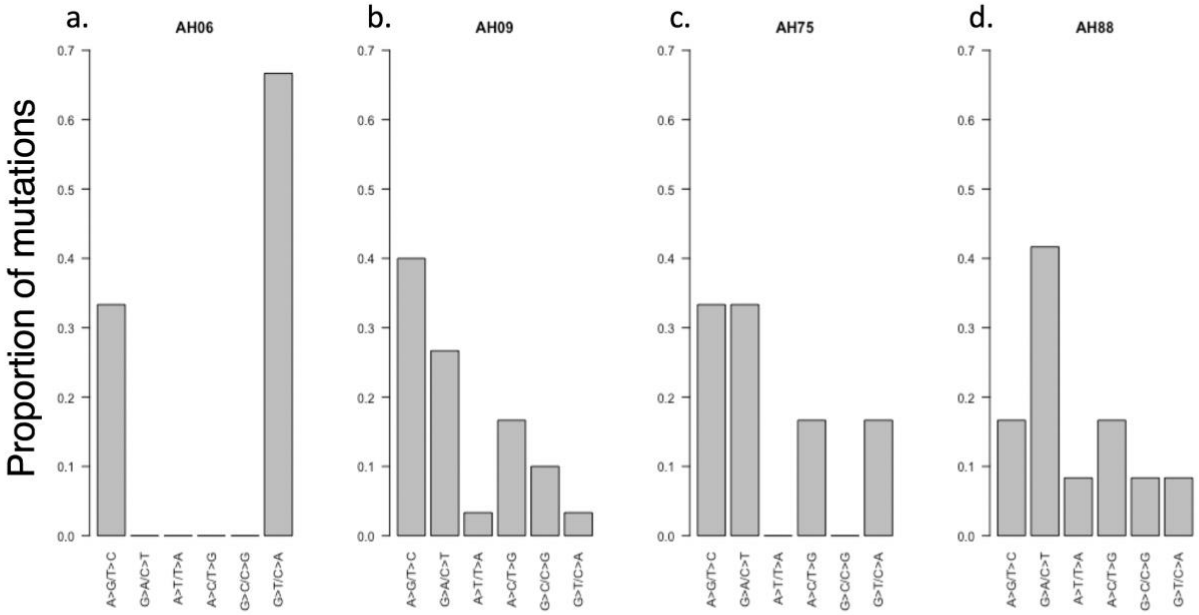


Figure S7. Mutation spectra for a.) verified AH06 mutations (N=3), b.) verified AH09 mutation (N=30), c. verified AH75 mutations (N=6), d.) verified AH88 mutations (N=12).

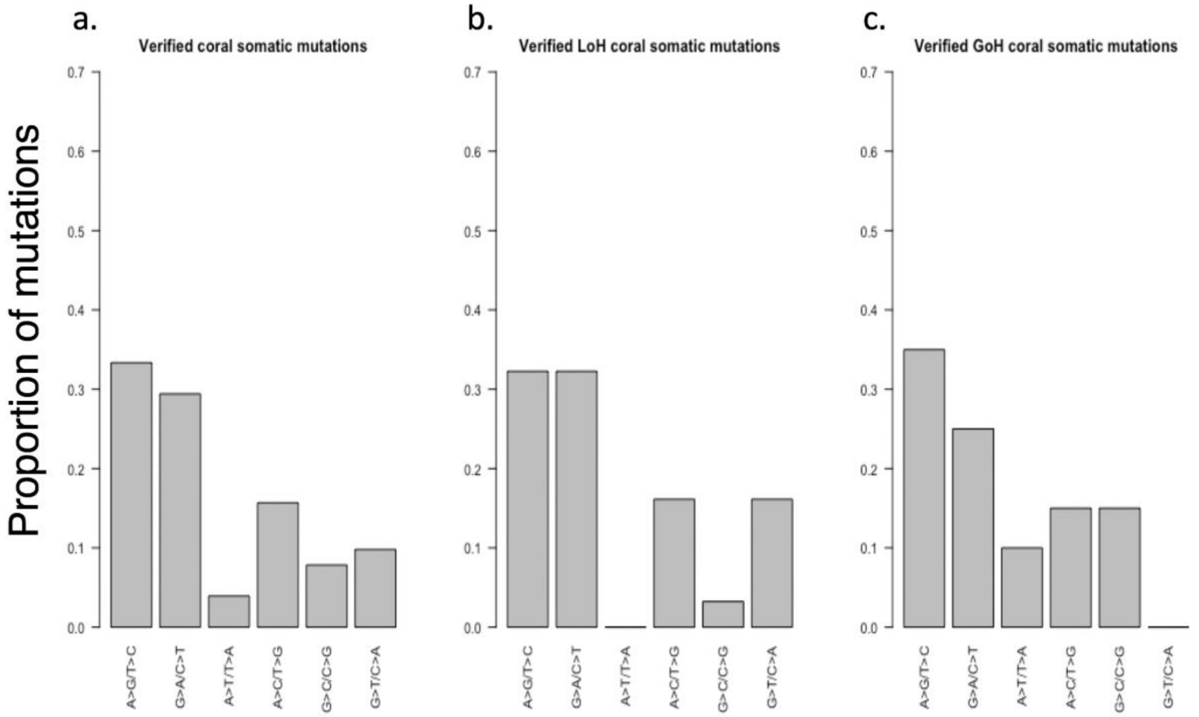


Figure S8. Mutation spectra for a.) all verified somatic mutations b.) all LoH somatic mutations, and c.) all GoH somatic mutations.

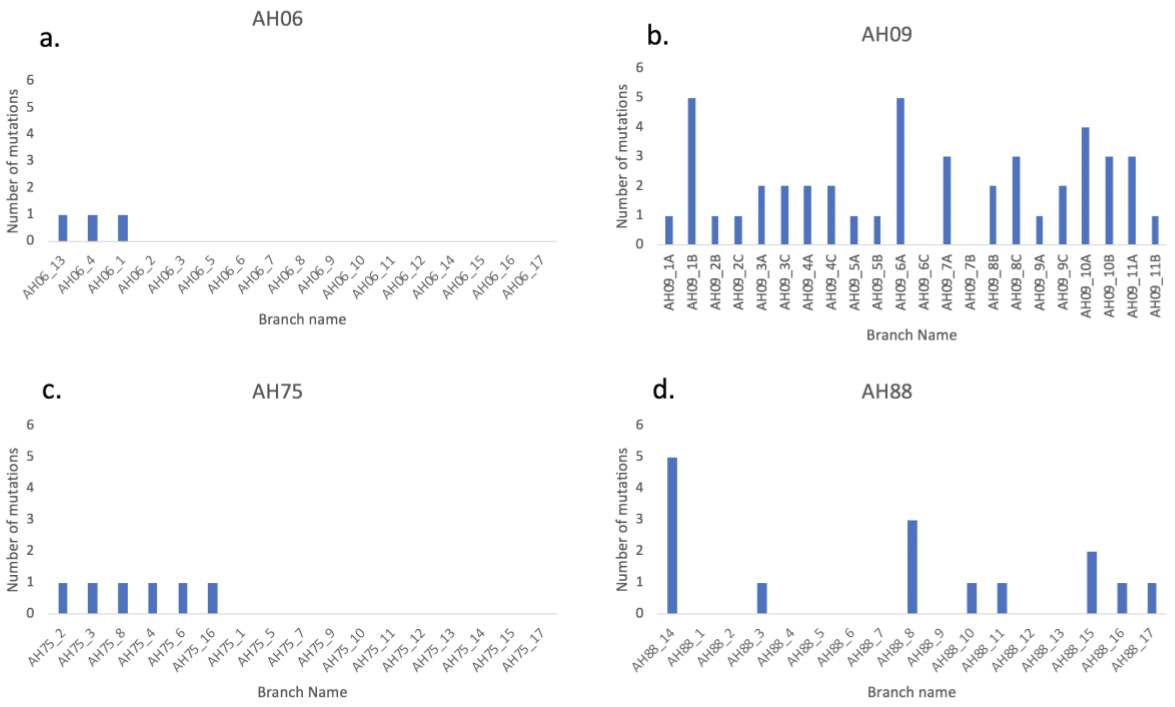

Figure S9. The number of mutations found in each branch of a colony for a.) AH06, b.) AH09, c.) AH75, and d.) AH88. Some mutations appeared in more than one branch. Branches AH09_1B and AH88_14 both had multiple linked mutation appear on a single colony (see Figure 5). In this figure, each of those linked mutations is treated independently. If they were treated as a single event, then AH09_1B would have 3 mutations instead of 5 and AH88_14 would have 2 mutations instead of 5.

**Supplemental tables**

Supplemental table S1. All putative somatic mutation data for AH06 (.txt file)

Supplemental table S2. All putative somatic mutation data for AH09 (.txt file)

Supplemental table S3. All putative somatic mutation data for AH75 (.txt file)

Supplemental table S4. All putative somatic mutation data for AH88 (.txt file)

Supplemental table S5. List of filtered, verified mutations for all colonies (.txt file)

Supplemental table S6. Estimates from CODEML analysis for colony AH88

| Model | Parameter estimates | dN/ds | lnL | Positively selected sites |
|---|---|---|---|---|
| M0 | 2.39 | 2.39 | -1457.2315 | none |
| M1 | p: 0.00001 0.99999<br>w: 0.06282 1.00000 | 1 | -1457.6283 | NA |
| M2 | p: 0.94040 0.00003 0.05957<br>w: 0.00000 1.00000 41.98518 | 2.5011 | -1455.2229 | **11, 29, 44, 100, 159, 216 (all p<0.01)** |
| M3 | p: 0.94043 0.00000 0.05957<br>w: 0.00000 0.00000 41.96817 | 2.5002 | -1455.2229 | **11, 29, 44, 100, 159, 216 (all p<0.01)** |
| M7 | p = 1.70896<br>q = 0.00500 | 1 | -1457.6283 | none |
| M8 | p0 = 0.94042<br>p = 0.00500<br>q = 1.43281, (p1 = 0.05958)<br>w = 41.96953 | 2.5003 | -1455.2229 | **11, 29, 44, 100, 159, 216 (all p<0.01)** |

Supplemental table S7. Estimates from CODEML analysis for colony AH09

| Model | Parameter estimates | dN/ds | lnL | Positively selected sites |
|---|---|---|---|---|
| M0 | w=dnds | 1.05638 | -7403.3188 | none |

| | | | | |
|---|---|---|---|---|
| M1 | p: 0.50001 0.49999 w: 0.15537 1.00000 | 0.5777 | -7403.0717 | NA |
| M2 | p: 0.68361 0.17318 0.14321 w: 0.46115 1.00000 1.24438 | 0.6666 | -7403.1704 | 168 G (nonsignficant) |
| M3 | p: 0.00001 0.99878 0.00121, w: 0.00000 0.73497 260.63945 | 1.0502 | -7396.2003 | **168 G (p<0.01)** |
| M7 | p = 0.00995 q = 0.00703 | 0.6 | -7402.9092 | none |
| M8 | p0 = 0.99877 p = 0.08440 q = 0.02933, (p1 = 0.00123) w = 260.41085 | 1.0586 | -7396.2069 | **168 G (p<0.01)** |

Supplemental table S8. Estimates from CODEML analysis for colony AH75

| Model | Parameter estimates | dN/ds | lnL | Positively selected sites |
|---|---|---|---|---|
| M0 | w= dN/ds | 0.493 | -1365.4664 | none |
| M1 | p: 0.99999 0.00001 w: 0.49296 1.00000 | 0.493 | -1365.4664 | NA |
| M2 | p: 1.00000 0.00000 0.00000 w: 0.49301 1.00000 1.00000 | 0.493 | -1365.4664 | none |
| M3 | p: 0.31376 0.49060 0.19564 w: 0.49294 0.49295 0.49306 | 0.493 | -1365.4664 | none |
| M7 | p = 97.03473 q = 99.00000 | 0.495 | -1365.4664 | none |
| M8 | p0 = 0.99999 p = 96.15305 q = 99.00000, (p1 = 0.00001) w = 1.00000 | 0.4927 | -1365.4664 | none |

Supplemental table S9. Estimated somatic mutation rate per gigabase per year

| **Colony** | **# of mutations** | **(# of gigabases assayed) * (# of samples)** | **Age in years (order of magnitude estimate)** | **Somatic mutation rate per gigabase per year** |
|---|---|---|---|---|
| AH06 | 3 | 0.04480714 | 10 | 6.7 |

| | | | | |
|---|---|---|---|---|
| AH09 | 30 | 0.11628780 | 100 | 2.6 |
| AH75 | 6 | 0.11024597 | 10 | 5.4 |
